# Effects of Substrate Stiffness on Neutrophil Adhesion over L-selectin Coated Endothelial

**DOI:** 10.1101/791434

**Authors:** Claude Guillory, Alice O. Dufour

## Abstract

Rolling of a cell under a hydrodynamic flow like the blood flow and the mechanism for the adhesion of a cell to the blood vessel is one of the fundamental process in many pathological and biological processes. An important example of these processes is inflammatory response and moving of the leukocytes to the sites of inflammation. While the blood-borne cells travel with the blood flow, they can interact with the inner endothelium’s wall, which is composed of a soft layer of endothelial cells. Not until recently, the effect of endothelial stiffness was poorly understood. Recent in-vitro and computational models, like modified Adhesive Dynamics, have shown that the elasticity of the underlying substrate can alter the rolling and adhesion of a cell. In this study, we investigate the effects of the substrate stiffness on the rolling and adhesion of a cell with neutrophil ligands by using the Adhesive Dynamic simulation. The vessel is modeled as an elastic surface coated with L-selectin molecules, which can form bonds with the ligands. In our simulation, the Young modulus of the surface ranges between 5 to 80 kPa. The results show that the softer substrate helps to capture the cell with neutrophil ligands. These results help us to understand how the state of adhesion changes for the neutrophil adhesion over L-selectin.

## 1. Introduction

Rolling and adhesion of a cell to a surface or another cell is a fundamental process in most of the biological process in a living being. Cells can travel through a living body by blood flow and attach to different surface based on the chemical binding characteristics. This process starts with forming bonds between binding molecules on the cell and the surface. Few examples of this process are, cancer metastasis, white blood cells attacking foreign agents in the body, and tissue regeneration [1-4]. This process is mediated by ligands as adhesion molecules on the cell and receptors as adhesion molecules on the substrate. The best-known ligands are neutrophils and PSGL-1 and the best examples for the receptors are selectins, cadherins, and integrins [5]. The most important receptor family is selectin family, which is consisted of endothelial cells (E-selectin), lymphocytes (L-selectin), and platelets (P-selectin) [6].

One of the important examples regarding the cell rolling and adhesion is the leukocyte accumulation in inflammatory response, known as the leukocyte-endothelial interaction. Endothelial layers are covered with L-selectin receptors, which mediate the capture of leukocyte with neutrophil or PSGL-1 ligands [7-9]. Part of this process is also leukocyte-leukocyte interaction, which is L-selectin-dependent [4-6 same], when leukocyte roll over a layer of other leukocytes attached to the activated endothelial cells. L-selectin is also known to be related to some diseases like diabetes, sepsis, and meningitis [10-15]. L-selectins also guide lymphocytes to find lymph nodes [16, 17]. In these examples, neutrophil-L-selectin binding is a vital process. Bond formation and rupture between neutrophil and L-selectin is a dynamic and stochastic process and depend on bond dissociation (*k*_*r*_) and formation (*k*_*f*_) rates, which themselves depend on the developed force in the bond. The recent studies have shown that the substrate elasticity play an important role in the state of adhesion of a rolling cell in a blood flow. Older experimental and theoretical experiment neglected these effects; however, the substrate elasticity can have an unknown impact in a specific cell adhesion. Also, in most of these studies the fluid is considered to be Newtonian, but recent studies on non-Newtonian fluids illustrate that those assumptions may not be perfect in particle movement or adhesion [18-21]. Computational modeling is a powerful tool to predict the behavior of a living organism and reduce the cost of the experimental studies. One of the important computational models to study the dynamic behavior of the cell rolling and adhesion is Adhesive Dynamic or AD, which was first developed by Hammer and Apte [22]. This model simulates the rolling and adhesion of a random cell over a random substrate and can be modified for any specific cell-surface interaction. Inspired by a recent experimental study [23], Moshaei et al. [2] modified AD algorithm to shows the effects of the surface elasticity on the adhesion over P and E-selectins covered surfaces. The authors used two-spring model first developed by [24] to add the surface elasticity as a parameter in the AD algorithm.

In the present study we used the same model as [2] to investigate the same effects for neutrophil rolling and adhesion over the L-selectin covered surface.

## 2. Methodology

In our simulation we used the model developed by Moshaei et al. [1, 2]. This model is based on AD simulation, first developed by Hammer and Apte in 1992 [22]. Moshaei et al. [2] added the substrate stiffness to the model in order to consider the effects of underlying substrate elasticity. As shown in the figure 1, the cell is a rigid spherical object with radius R in a viscous shear flow with μ and 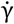 as its viscosity and shear rate, respectively. The surface is coated with L-selectin receptors and is modeled as an incompressible elastic half space with Young modulus E and ν = 0.5for the Poisson’s ratio.

**Figure 1.**
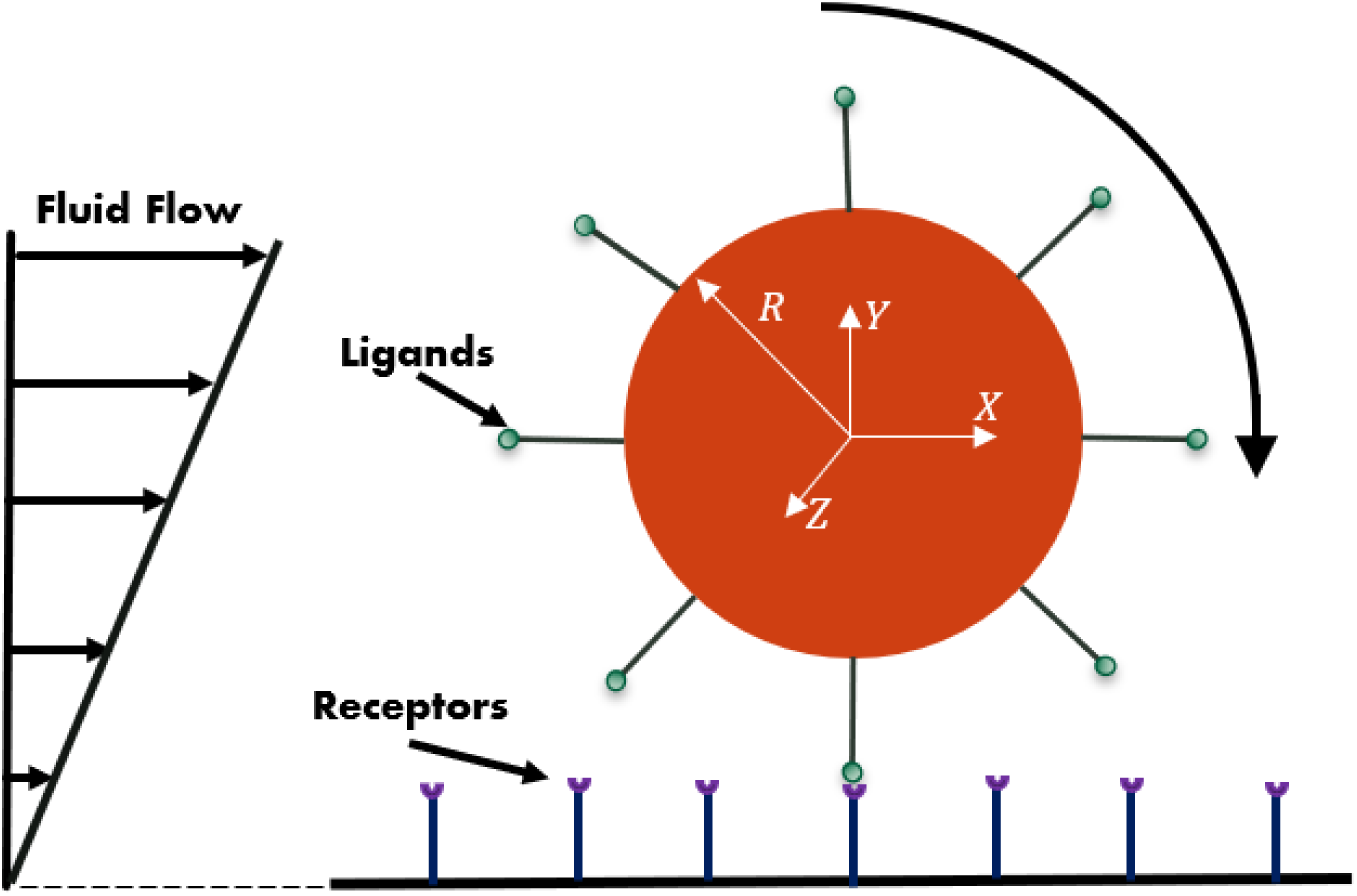
Rolling cell subjected to shear flow force and torque.

AD simulation uses the Monte Carlo stochastic algorithm to update the cell’s velocity in each computational time step. After all the forces on the cell are known at each time step, the simulation uses the Monte Carlo probability functions and updates the number of free ligand and receptors as well as the number of the bonds between the cell and the substrate. Based on the original model [22], the probabilities of bond formation P_f_ and rupture P_r_ are,

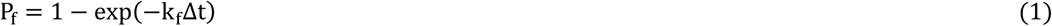

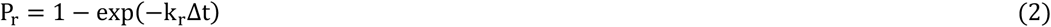

In the above equations, k_f_ and k_r_ are bond formation and rupture rates, respectively. Bell [25], formulated the reverse reaction rate k_r_ based on the developed force in the bond between the ligand and the receptor,

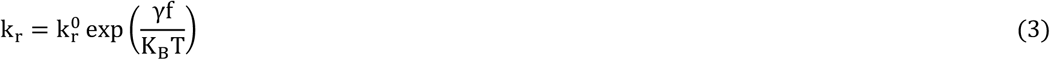

Here, K_B_ is the Boltzmann constant, T is the absolute temperature, 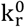 is constant reverse rate, γ is the reactive compliance, and f is the developed force in the bond.

The bond formation rate in this model is,

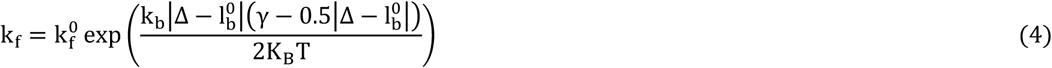

Here, 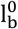 is the equilibrium bond length, k_b_ is the equivalent bond spring constant, 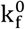 is the constant forward rate, and Δ is the distance between the bonding molecules.

As shown in figure 2, the stiffness of the substrate is considered to behave as an elastic spring. Following Kendall [26] model, the stiffness can be expressed as,

**Figure 2.**
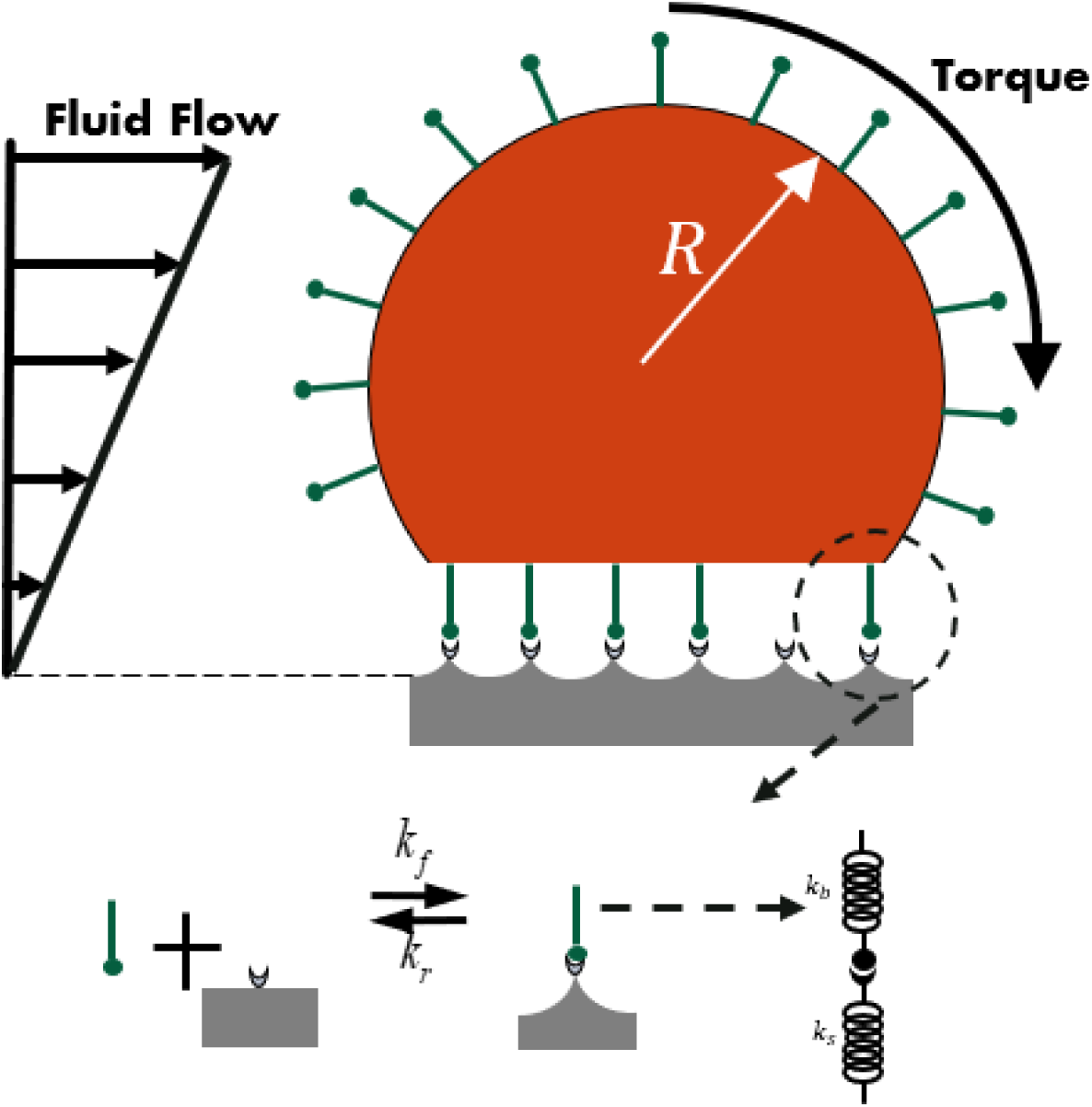
Bond formation between the cell and the substrate and two-spring model.

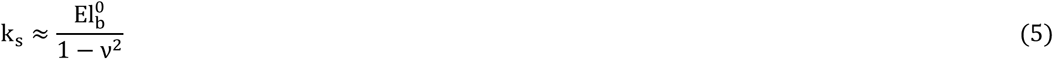

while l_b_ is the length of the bond, the developed force in the bond is,

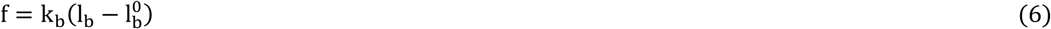

So, the total equivalent stiffness of the bond and the surface is,

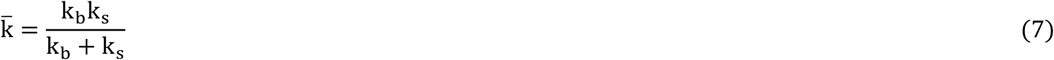

The values for the AD simulation are presented in table 1.

**Table 1.** Simulation parameters and values [27, 28].

| Constant | Definition | Value |
| --- | --- | --- |
| R | Cell radius | 5 $\mu\text{m}$ |
| $k_b$ | Bond stiffness | 1 dyn/cm |
| $l_b^0$ | Equilibrium bond length | 25 nm |
| $\Delta t$ | Time step | $10^{-6}$ sec |
| $\mu$ | Viscosity | 1 g/cm. s |
| $\dot{\gamma}$ | Shear rate | $100 \text{ s}^{-1}$ |
| $\gamma$ | Reactive compliance | 1.11 $\text{\AA}^0$ |
| $N_l$ | Ligand density | 3500 |
| $k_f^0$ | Intrinsic forward rate | $84 \text{ sec}^{-1}$ |
| $k_r^0$ | Intrinsic reverse rate | $2.8 \text{ sec}^{-1}$ |

## 3. Results and Discussions

For our simulations we used the same values for the parameters as used in the previous studies [1, 2, 3, 22, 27, 28]. The Young modulus for the surface elasticity ranges between 5 to 80 kPa and to better illustrate the results, we only present the results for 5, 20, and 80 kPa. The constant forward and reverse rates for bond formation and rupture between neutrophil and L-selectin is provided in table 1.

Figure 3 illustrates how the total number of bonds varies at each step. This is due to the stochastic nature of the bond formation and rupture between the cell and the surface. The total number at each time step is plotted for all three different elasticity together. As it is clear in the figure, the total number of the bonds and its fluctuations is smaller as the surface become comparatively more rigid. This means that at each time, the are less bond and force to hinder the cell’s rolling. In the other words, the bonds can be stretched more when they are interacting with a softer substrate and resist against shear force and torque.

**Figure 3.**
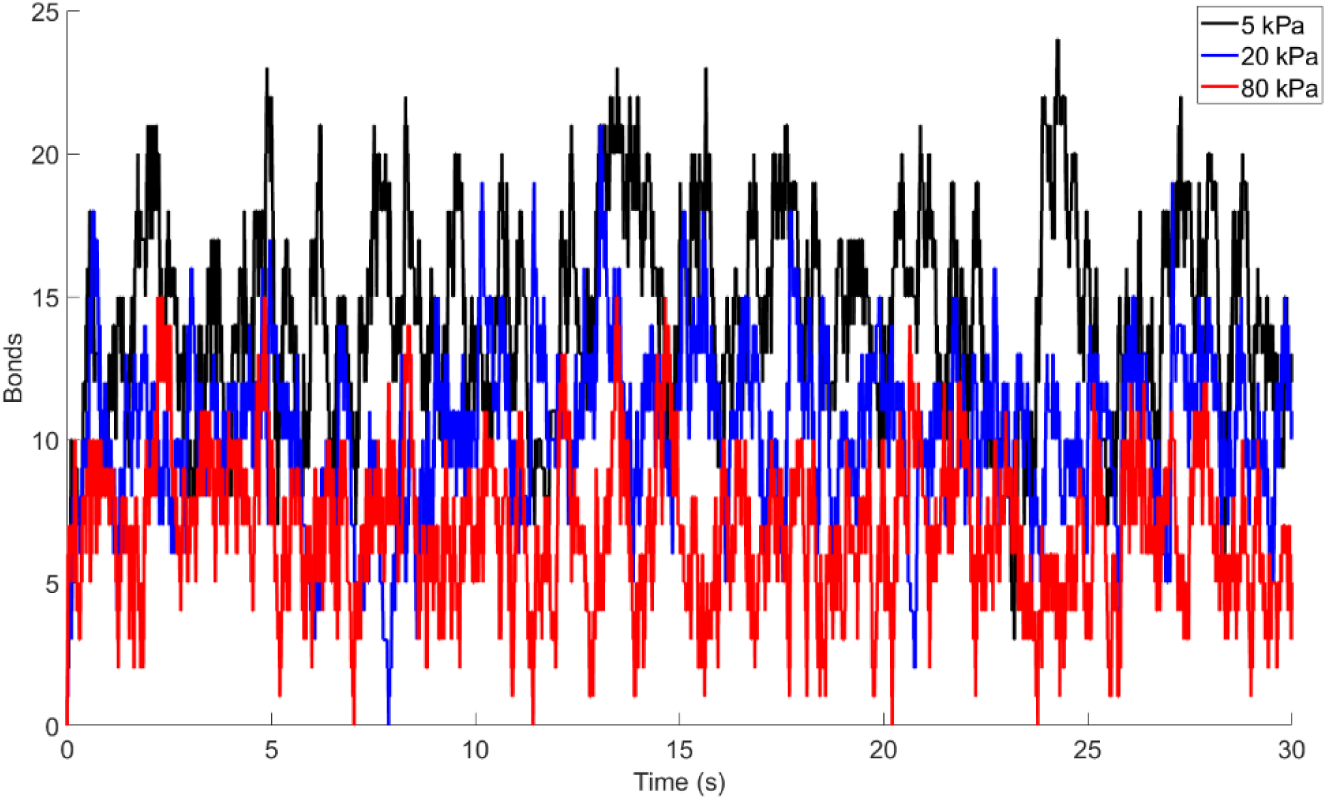
Fluctuations of the total number of bonds for a rolling cell over surfaces with E = 5, 20, 80 kPa.

The average number of bonds are 7, 12, 16, for 5, 20, and 80 kPa, respectively.

The most interesting result of this study is the effects of the surface elastic on the cell’s velocity and the distance that cell can travel. Figure 7 shows the comparison for the surface with different stiffness. As it can be seen in this figure, as the surface stiffness decreases, the developed force in the bonds can resist more against the fluid forces. This figure shows that the cell travelled more over the stiff substrate rather than a surface with E = 5 kPa. These results are totally different than what previously were presented for P and E-selectins, in computational [1, 2] and experimental studies [16]. One common conclusion in most of the recent studies is that the surface’s Young modulus is not the only factor in the cell rolling and adhesion. There are many different parameters that can change the rolling and adhesion behavior and the surface elastic is only one of them. However, the observed trend here is new, in none of the previous studies the cell could travel more over the rigid substrate rather than the softer one. Couple of the parameter that may have an important role in this trend are the forward and revers reaction rates as well as the reactive compliance.

**Figure 4.**
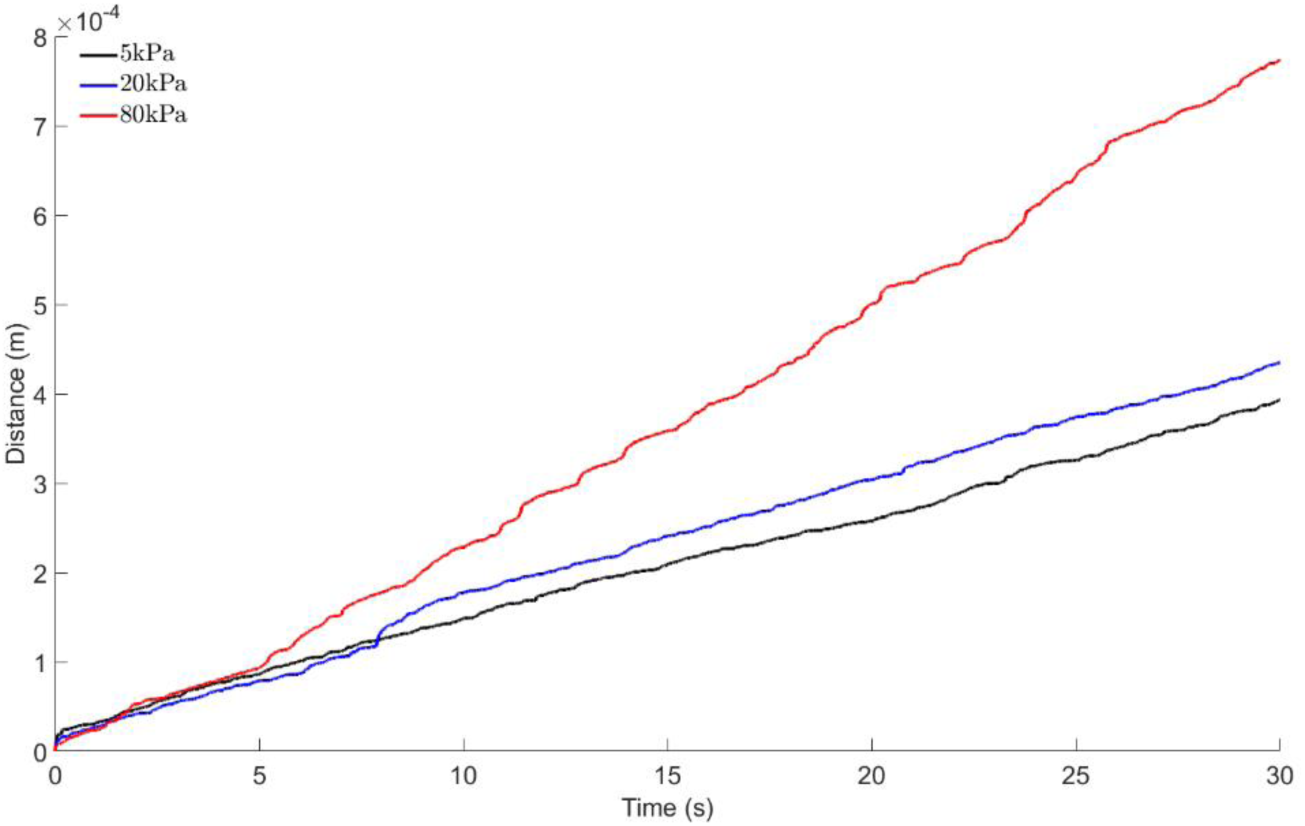
Travelled distance of the cell while formed bonds resist the sear flow.

**Figure 5.**
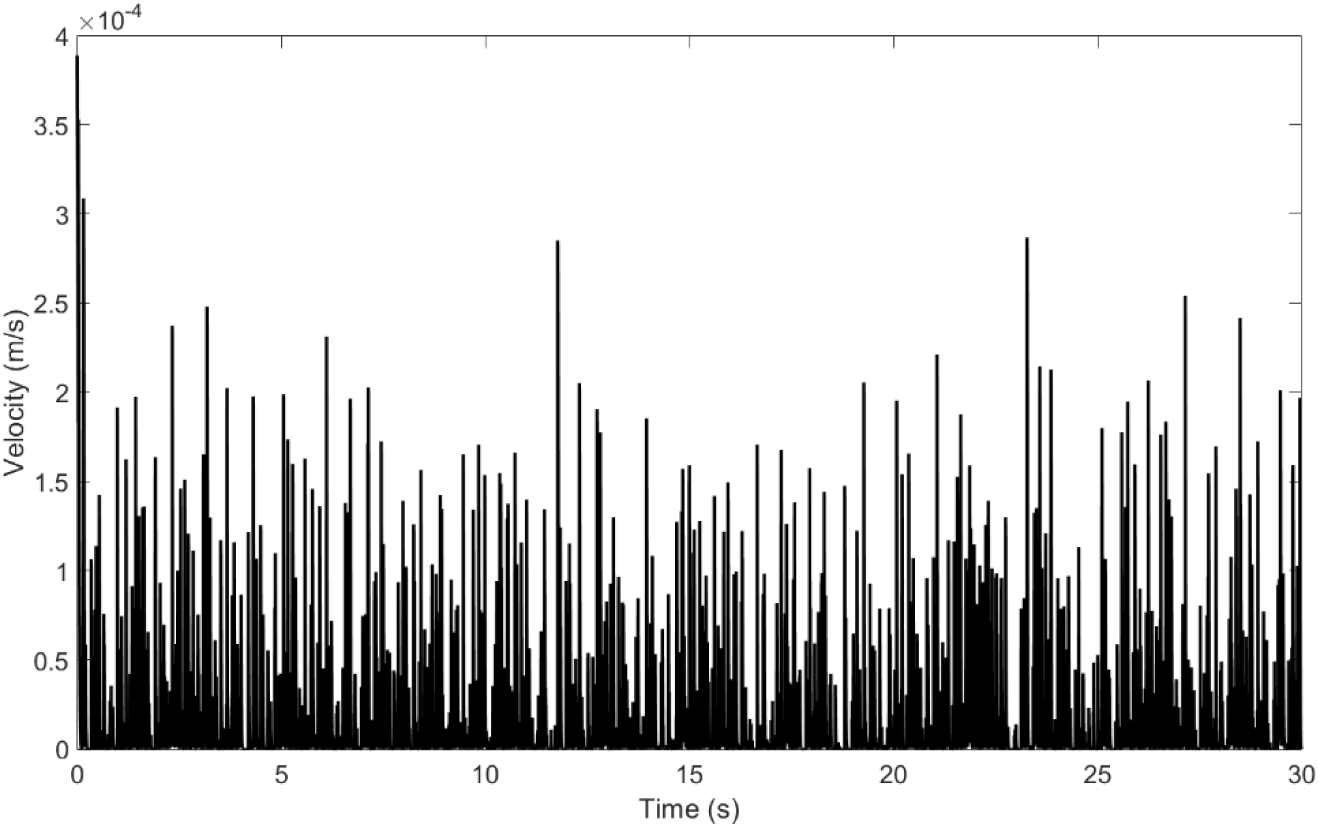
Velocity distribution of the cell over surface with E=*5* kPa.

**Figure 6.**
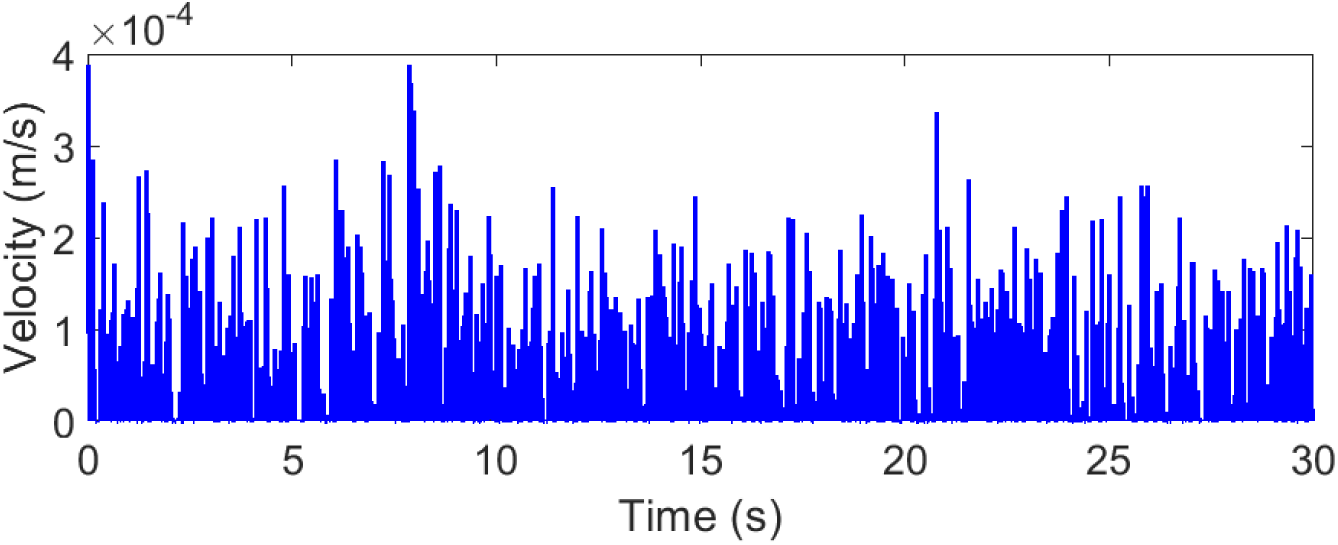
Velocity distribution of the cell over surface with E=20 kPa.

**Figure 7.**
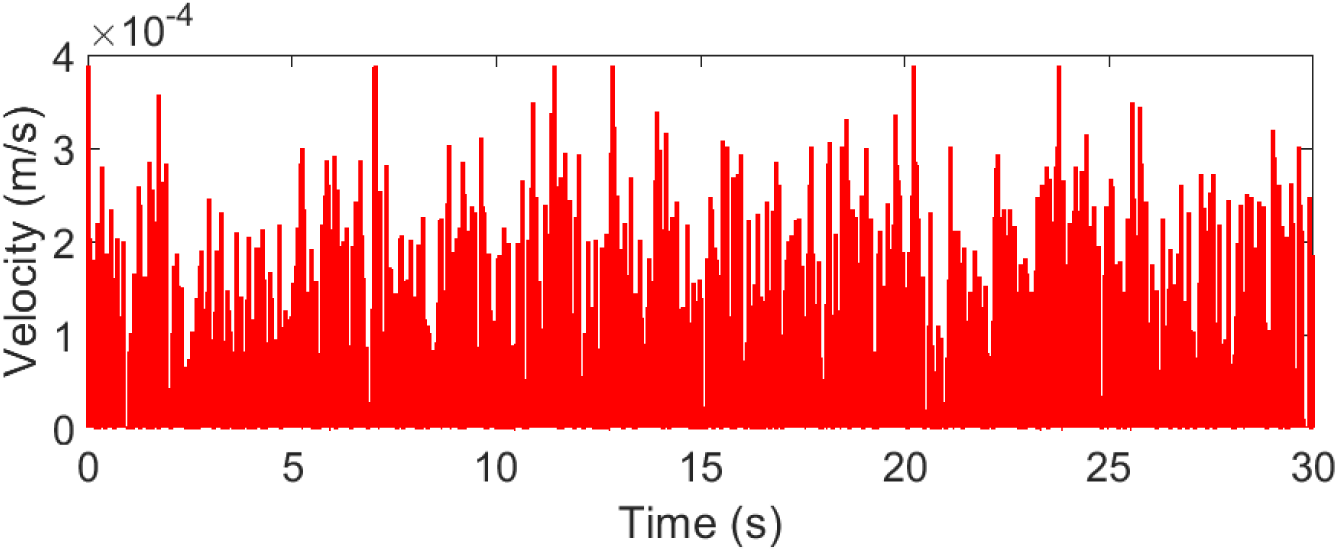
Velocity distribution of the cell over surface with E=80 kPa.

Figures *5*, 6, and 7 illustrate the changes in the velocity of the cell in 30 seconds of the simulations. The important result here is the scale of the fluctuations and the magnitude of the average velocity over stiffer surface. As it can be concluded from these figures, when the cell rolls over the surface with E = 80 kPA the fluctuations of the instantaneous velocity is bigger and the average velocity is 1.4 m/s, which is consistent with the conclusion from figures 3 and 4, and means there are less resistive forces against the shear flow, so the cell could travel more and survive the capture by the surface.

## 4. Conclusion

This study focuses on the neutrophil and L-selectin adhesion for a rolling cell in a shear flow like blood flow. We investigated the effects of the surface rigidity by employing the AD simulation and changing the elasticity of the surface using two-spring model. Although we used the experimental data for our parameter, yet currently, there is no experimental studies to check our predicted results. In this study we use three different Young moduli 5, 20, and 80 kPa.

the results of this study showed that the effects of the surface elasticity are different than what were shown for P and E-selectin in previous studies [1, 2, 16]. In those studies, the surface elasticity has a minimal effect on the rolling over P-selectin coated surface, while it can change the state of adhesion for the rolling over E-selectin coated surface. Here, the effect is totally different. Our model predicts that the rigidity does not help to capture the cell and on the other hand, as the surface elasticity increases the chances are more that the surface captures the cell. As mentioned in the result section, there are many different parameters that can change the state of adhesion and the surface elasticity is only one of them. An important conclusion here is that it is not possible to predict the effects of surface elastic based on only few models and a greater number of simulations with ability to change different parameters are needed to be able to come to a more comprehensive conclusion of the general cell rolling and adhesion.

## Appendix

In this section we discuss the forces on the cell and how these forces are calculated in AD simulation. In this model, the cell is small and moves with the flow so, the inertia effects are neglected, and the equilibrium is written as,

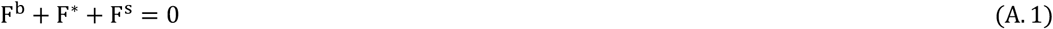

Where, F^s^ is the shear force, F*is the resultant non-specific forces, and F^b^ is the force in the all bonds. Non-specific forces are, the electrostatic 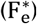, the Van der Walls 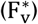, the gravity 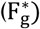, and the steric stabilization forces 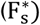 and are defined below (the parameters and their values are defined in table 2).

**Table 2.** Non-specific forces parameters and values [1, 2].

| Constant | Definition | Value |
| --- | --- | --- |
| $A_h$ | Hamaker constant | $5 \times 10^{-21} \text{ J}$ |
| $\rho_f$ | Fluid density | $1.05 \text{ g/cm}^3$ |
| $L$ | Cell coat thickness | $10 \text{ nm}$ |
| $\rho$ | Charge density | $0.05 \text{ g/cm}^3$ |
| $\rho_c$ | Cell density | $1 \text{ g/cm}^3$ |
| $H$ | Surface thickness | $70 \text{ \AA}^0$ |
| $\kappa$ | Debye-Huckel length | $0.125 \text{ \AA}^0$ |
| $\epsilon$ | Dielectric constant | $7.8 \times 10^{-19} \text{ C}^2/\text{dyn. cm}^2$ |
| $\lambda$ | Steric constant | $2.5 \times 10^{-6} \text{ dyn}$ |

Electrostatic:

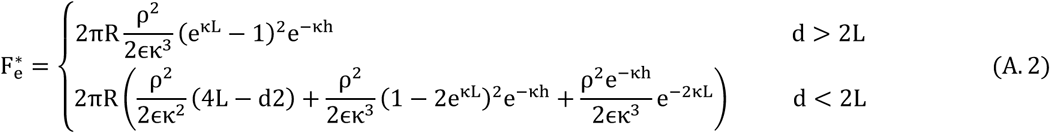

Van der Walls:

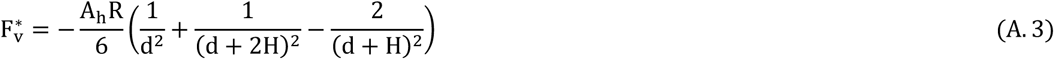

Gravity:

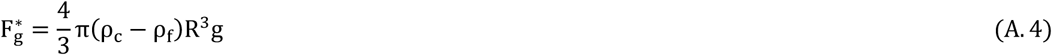

Steric stabilization:

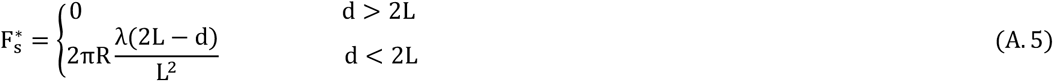

In order to calculate share force F^h^ and shear torque C^h^, Goldman et al. [29, 30] were followed,

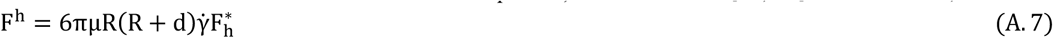

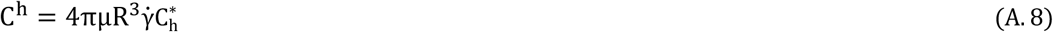

Here, 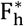 and 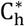 are given by [29, 30] and depend on the distance between the cell and the surface.

To calculate the velocity at each time step Mobility matrix is used at the equilibrium condition as

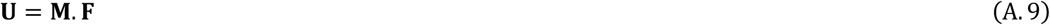

**M** is a 6 × 6 mobility matrix defined by [29-31],

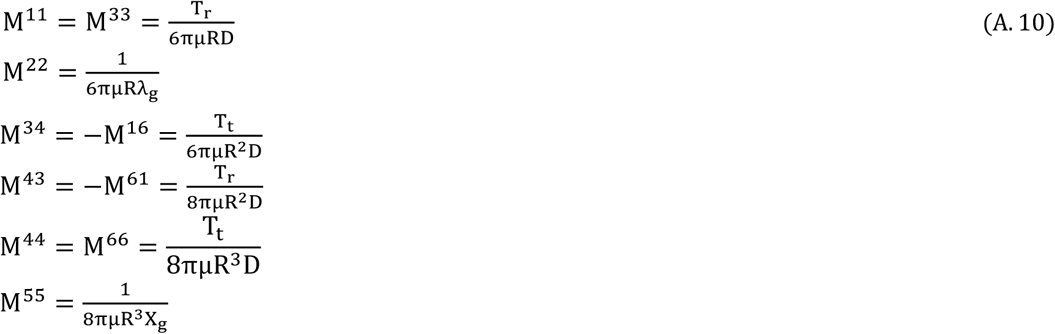

***U*** is the velocity vector,

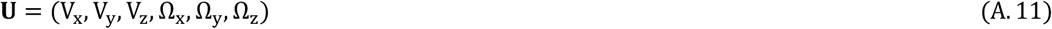

**F** is the force vector,

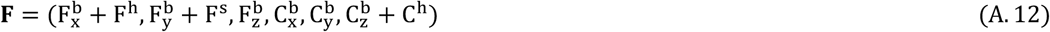

V and Ω are the linear velocity and the angular velocity, respectively.

